# Absence of homeostatic downscaling in dentate gyrus granule cells

**DOI:** 10.64898/2026.02.13.705706

**Authors:** Owen D. Jones, Kayla Benjamin, Ruth M. Empson, Wickliffe C. Abraham

## Abstract

Granule cells of the hippocampal dentate gyrus fire at exceptionally low rates in order to maintain sparse neural codes. Despite the need for granule cells to remain in such relative quiescence, the mechanisms that regulate their firing rates have not received much attention. We investigated the potential of family of mechanisms, homeostatic downscaling, that could theoretically maintain granule cell firing at preferential levels following chronically elevated activity. Surprisingly, we found no evidence of reduced synaptic input or intrinsic excitability in granule cells even after prolonged exposure to GABA_A_ receptor blockade. In fact, we found that mini excitatory postsynaptic current frequency was elevated in granule cells after prolonged exposure to GABA_A_ antagonists. This effect was consistent across blockers or when cell firing was driven by elevated extracellular K^+^, and did not rely on NMDA receptors, L-type voltage gated Ca^2+^ channels or thrombospondin-driven synaptogenesis. However, the magnitude of long-term potentiation was reduced at synapses onto granule cells after prolonged exposure to a GABA_A_ antagonist *in vivo*. We conclude that granule cells are the first known cell type that do not display homeostatic downscaling. Instead, these cells rely on other mechanisms, including metaplasticity, to maintain their activity within optimal bounds.

## Introduction

Regulation of neuronal activity is critical for healthy brain function, as attested by the number of neuropathologies characterised by aberrant activity in neural circuits (Bakker et al., 2012; Ellwardt et al., 2018; Huijbers et al., 2019; Yizhar et al., 2011). Activity is regulated by a plethora of mechanisms, among them “homeostatic plasticity” (Turrigiano, 2011; Turrigiano, Leslie, Desai, Rutherford, & Nelson, 1998), in which chronic firing rate perturbations are countered by negative feedback mechanisms acting on synapse weight and intrinsic excitability, and “metaplasticity” (Abraham, 2008; Hulme, Jones, & Abraham, 2013), which entails the activity-dependent regulation of future neural plasticity. Such mechanisms have been demonstrated in various regions of cortex and in the hippocampal CA subfields, and are considered critical for network stability and healthy cognition (Echegoyen, Neu, Graber, & Soltesz, 2007; Philpot, Espinosa, & Bear, 2003; Singh, Sateesh, Jones, & Abraham, 2022; Wen, Prada, & Turrigiano, 2025; C. H. Wu, Ramos, Katz, & Turrigiano, 2021).

The hippocampal dentate gyrus (DG) is a region that, in theory, requires exceptionally tight control of activity in order to function properly. As the gateway for the majority of cortical input to the hippocampal formation, the DG receives projections from layer II entorhinal cortex (EC) pyramidal cells that are sparsely distributed across the considerably greater granule cell (GC) population (Amaral, Ishizuka, & Claiborne, 1990; Amaral, Scharfman, & Lavenex, 2007). This decorrelation of cortical information is maintained by the exceptionally low firing rates of GCs (Diamantaki, Frey, Berens, Preston-Ferrer, & Burgalossi, 2016; Pernía-Andrade & Jonas, 2014; Senzai & Buzsáki, 2017), which enable segregated streams of information to be conveyed downstream to the CA3 region for continued segregation or recombination as required (Leutgeb, Leutgeb, Moser, & Moser, 2007). Yet, despite the criticality of sparse GC firing, few studies to date have investigated the plasticity mechanisms that maintain low levels of GC activity.

Here, we used organotypic slices of hippocampus and EC to probe the consequences of chronic activity elevation on GCs. In contrast to other glutamatergic cell types of the hippocampus (Goold & Nicoll, 2010; Lee et al., 2013; Mao et al., 2018), we found no evidence of homeostatic downregulation of evoked firing or synaptic activity in GCs following chronically elevated activity. Instead, GCs respond to prolonged GABAergic blockade with *enhanced* rather than reduced synaptic transmission. Specifically, we found that 120 hr (5 days) of GABAergic blockade triggers an increase in miniature excitatory postsynaptic current (mEPSC) frequency. The effect was visible across a range of treatments aimed at boosting activity, was not due to competing “Hebbian” plasticity driven by NMDARs, did not rely on thrombospondin-mediated synaptogenesis, and was not seen in cells of the neighbouring CA3 region. However, we found that prolonged treatment with a GABAergic antagonist *in vivo*, while exerting no overt effects on synaptic transmission, metaplastically reduced subsequent long-term potentiation at EC-GC synapses. We conclude that homeostatic synaptic downregulation (termed “downscaling” elsewhere) is absent in GCs, which instead rely in part on metaplasticity to maintain activity within preferential limits.

## Methods

### Organotypic slice culture preparation

All protocols were conducted with approval from the University of Otago’s Animal Ethics Committee. Slice cultures were prepared from P4-6 C57Bl/6J mouse pups of either sex following validated methods that preserve the fundamental features of the entorhino-hippocampal circuitry (Del Turco & Deller, 2007; Radic et al., 2017). Briefly, mice were decapitated are brains dissected free into ice-cold Minimal Essential Media (MEM, Invitrogen) supplemented with 2 mM Glutamax, 20 mM HEPES, 0.45% (w/v) glucose, 100 U/ml penicillin and 0.1 mg/ml streptomycin (pH 7.3–7.4). Horizontal brain slices (350 µm) were prepared using a Vibratome (Leica), from which sections including hippocampus and associated EC were isolated under a dissection microscope. Sections were transferred to porous membrane inserts (Millicell PICM0RG50, 0.4 μm pore size) and maintained in 6 well plates at interface between atmosphere and 1 ml of culture media (42% MEM, 25% Basal Eagle Medium with Earle’s salts, 25% heat-inactivated horse serum, 0.65% glucose, 25 mM HEPES, 0.1 mg/ml streptomycin, 100 U/ml penicillin, 0.15% sodium bicarbonate, 2 mM Glutamax, pH 7.30 with NaOH). Cultures were maintained in a humidified incubator (5% CO_2_) at 34°C for 14 days prior to treatment, with media replaced every 2-3 days.

### Drug treatments

Beginning DIV14, cultures were treated with either standard (control) media or media supplemented with one of the following: 25 µM gabazine hydrobromide; 100 µM picrotoxin; 100 µM picrotoxin & 4 mM KCl; 100 µM picrotoxin, 4 mM KCl & 50 µM D-APV; 100 µM picrotoxin & 100 µM gabapentin. Picrotoxin was dissolved in DMSO. All other compounds were dissolved in H_2_O. Drugs were diluted 1000x from stock, and control cultures were treated with corresponding amounts of the appropriate vehicle. In the case of KCl treatment, equimolar NaCl was also added to control cultures to account for altered osmolality. All drugs were purchased from Hello Bio, except for gabapentin which was purchased from MedChemExpress.

### Patch clamp electrophysiology

After 1-20 days (24-480 hr) of treatment, slice cultures were transferred to a recording chamber attached to a stage and upright microscope (Olympus BX50WI) equipped with infrared differential interference contrast optics for visualising individual cells. Slice cultures were superfused with room temperature (24 °C) artificial cerebrospinal fluid (aCSF, 124 mM NaCl, 3.2 mM KCl, 1.25 mM NaH_2_PO_4_, 26 mM NaHCO_3_, 2.5 mM CaCl_2_, 1.3 mM MgCl_2_, and 10 mM d-glucose) saturated with 95% O2 / 5% CO2, flowing at 2 ml/min. The DG or CA3 region was identified under low (5x) magnification and individual GCs of the DG crest or pyramidal cells of the proximal CA3 region were visualised and targeted under 40x magnification. Cells were patched with thick-walled borosilicate glass recording pipettes pulled on a Flaming/Brown micropipette puller (Sutter P-97) giving tip resistances of 2-5 MΩ when filled with internal solution. Recordings were made via a MultiClamp 700B amplifier and Digidata 1440A A/D converter, using MultiClamp Commander and pClamp10 software (all hardware and software from Molecular Devices). Cells were patched and allowed to stabilise for 5 min before recording. Cells were kept for recording if resting V_m_ was greater than -70 mV for GCs or -60 mV for CA3 pyramidal cells immediately (< 10 s) after break-in. Cells were included in analyses if R_a_ remained < 20 MΩ and was < 10% of R_m_, and if neither parameter varied by > 20% over the course of the experiment.

*Current clamp recordings:* pipettes were filled with 130 mM K^+^gluconate, 10 mM HEPES, 4 mM ATP-Na, 0.4 mM GTP-Na_2_, 10 mM phosphocreatine, 4 mM MgCl_2_ (pH 7.3 with KOH, 295 mOsm). Signals were filtered at 10 KHz and digitized at 20 KHz. Holding current was manually applied to maintain resting V_m_ at -70 mV. R_in_ was calculated from the steady-state (i.e. post-voltage sag) V_m_ deflection in response to 750 ms hyperpolarising current steps (-20 pA) from 0 to -200 pA and dividing the former value by the latter at each step. Evoked firing was assessed as the number of action potentials (APs) triggered in response to 500 ms depolarising current steps (50 pA) from 0-200 pA.

In acute drug application studies, cell firing was assessed on DIV 14 using current clamp conditions as above, but with cells held at resting V_m_ while recording in gap-free mode. To best replicate culture conditions, cultures were held at 34°C.

*Voltage clamp recordings*: pipettes contained 90 mM KMeSO4, 40 mM CsMeSO4, 10 mM HEPES, 4 mM ATP-Na, 0.4 mM GTP-Na2, 10 mM phosphocreatine, 4 mM MgCl_2_, 0.2 mM EGTA and 4 mM QX-314 (pH 7.3 with CsOH, 295 mOsm). Cells were voltage clamped at -70 mV. To isolate AMPA receptor-mediated mEPSCs, the aCSF was supplemented with 1 µM TTX, 50 µM APV and 20 µM gabazine. 5 min epochs were recorded in gap-free mode with 2 KHz filtering and 10 KHz sampling. The liquid junction potential was not compensated for.

### Field potential recordings

Drug treatment and slice preparation: C57/B6 mice aged 2-3 months of either sex were maintained on a 12 hr light/dark cycle with food (chow) and water *ad libitum*. Mice were injected (i.p.) twice daily for 3 days with 1 mg/kg PTX freshly dissolved in 0.9% saline at 1µg/1µL (w/v). Control mice received saline-only injections.

Twelve hr post final injection, horizontal brain sections including hippocampus and EC were prepared as previously described (Jones, Hulme, & Abraham, 2013). Briefly, mice were deeply anesthetised with isoflurane inhalation and decapitated. Brains were rapidly removed and placed in ice-cold dissection solution (mM: 210 mM sucrose, 26 mM NaHCO3, 2.5 mM KCl, 1.25 mM NaH2PO4, 0.5 mM CaCl2, 3 mM MgCl2, 20 D-glucose) gassed with 95% O2/5% CO2. 400 µm sections were prepared using a Vibratome (VT1000, Leica) and were left to recover in a humidified chamber at the interface between air and aCSF. Slices remained at 32°C for 30 min post-sectioning, then at room temperature for at least 90 min prior to use.

For recordings, slices were transferred to a recording chamber and submerged in aCSF (32.5°C) at a flow rate of 2 ml/min. Synaptic field excitatory postsynaptic potentials (fEPSPs, > 2 mV amplitude at 0.1 mA) were evoked through stimulation of the medial performant path inputs to the DG using custom built stimulators coupled to 50 μm Teflon-insulated tungsten monopolar electrodes (diphasic pulse half-wave duration 0.1 ms) placed in the inner blade. Recordings were made using glass micropipettes filled with aCSF (2-3.5 MΩ) connected to Grass™ preamplifiers. Electrodes were placed centrally within the molecular layer approximately 400 μm apart. Correct recruitment of medial performant path axons was confirmed through paired-pulse tests (50 ms interpulse interval) that revealed paired-pulse depression of the fEPSP in all cases. There were no differences in baseline fEPSP slope or paired-pulse ratio between groups (fEPSP slope, *p* = 0.13; PPR, *p* = 0.45; data not shown). For LTP experiments, 0.2 µM gabazine was then added to aCSF. Baseline stimulation (2-3 mV amplitude at 65 μA) consisted of single pulses delivered every 30 s for 15 min. LTP was then induced via 4 trains of bursts (10 bursts at 5 Hz/train, 10 x 100 Hz pulses/burst, intertrain interval 30 s). Test pulses then resumed for 60 min post-LTP induction.

### Data analysis

All analysis was conducted blind to condition. Current clamp data were analysed using Easy Electrophysiology software with APs classified as events overshooting 0 mV and steady state V_m_ taken as the averaged V_m_ during the final 50 ms of the current step. Voltage clamp data were extracted and analysed with MiniAnalysis software by first using recommended detection parameters for AMPA receptor-mediated mEPSCs (> 5 pA amplitude), then visually inspecting each frame of the recording to check for false positives/negatives and accurate measurement of events. A sample of < 200 mEPSCs/cell (for GCs) or 150 mEPSCs/cell (CA3 neurons) was used to generate mean amplitudes, frequencies and kinetic measurements. Field potentials were analysed using custom software designed using LabView. The initial linear portion of the fEPSP slope was taken as a measure of synaptic efficacy. The averaged fEPSP slope for the final 10 min prior to LTP induction was taken as a baseline, and all measurements were then expressed as a percentage of this value. LTP was assessed as the averaged % change in fEPSP slope during the final 10 min of recording.

Statistical analyses of group means were made with GraphPad Prism 10.0, using Student’s *t* test (two tailed) or ANOVA with post-hoc Tukey test, as appropriate. A *p* value of < 0.05 was considered statistically significant.

## Results

### Chronic GABA_A_ blockade leads to augmented, not reduced synaptic transmission onto granule cells

GABAergic antagonists are the most common method used to augment neuronal activity and induce downward homeostatic plasticity of synapses (Siddoway, Hou, & Xia, 2014). Thus, we began probing for this form of plasticity in granule cells using a saturating concentration (25 µM) of the GABA_A_ receptor antagonist gabazine. Curiously, voltage clamp experiments revealed no effect of gabazine on mEPSC amplitude or frequency with < 72 hr of gabazine treatment, compared to controls. Further, after 120 hr of gabazine treatment, we not only failed to see evidence of synapse downregulation, but instead saw no change in mEPSC amplitude and an *increase* in mEPSC frequency (**Fig. 1**). No effects of treatment on mEPSC kinetics (rise/decay time) were observed at any timepoint. Current clamp experiments revealed only a modest reduction in input resistance (R_in_) after 24 or 120 hr of gabazine treatment, which did not translate to any discernible difference in excitability as assessed via evoked firing. Resting membrane potential (RMP) was also unchanged by gabazine treatment at either timepoint (**Fig. 2**)

**Figure 1:**
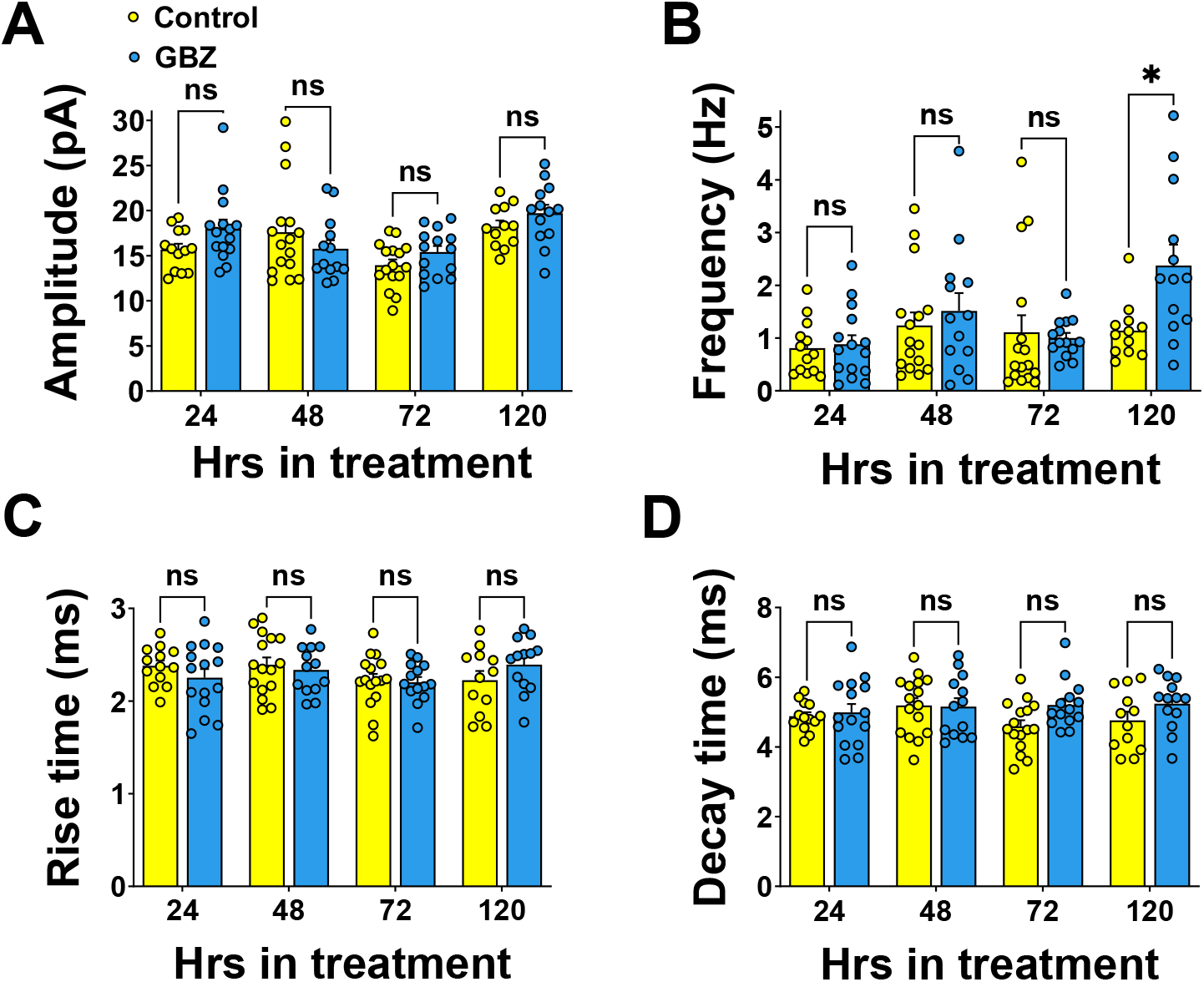
Chronic gabazine treatment augments synaptic transmission onto dentate gyrus granule cells. **A**: Pairwise comparisons of control vs. treated cells after 24, 48, 72 and 120 hrs of treatment reveal no significant differences in mEPSC amplitude. **B**: gabazine elevates mEPSC frequency after 120 hrs of treatment (Control: 1.14 ± 0.2 Hz; GBZ: 2.38 ± 0.4 Hz, *p* = 0.037). **C** and **D**: no effect of treatment on mEPSC kinetics at any timepoint. (n/group: 24hr: Control = 13, Gabazine = 15; 48 hr: Control = 16, Gabazine = 13; 72 hr: Control = 16, Gabazine = 14; 120 hr: Control, n = 12; Gabazine, n = 13)

**Figure 2.**
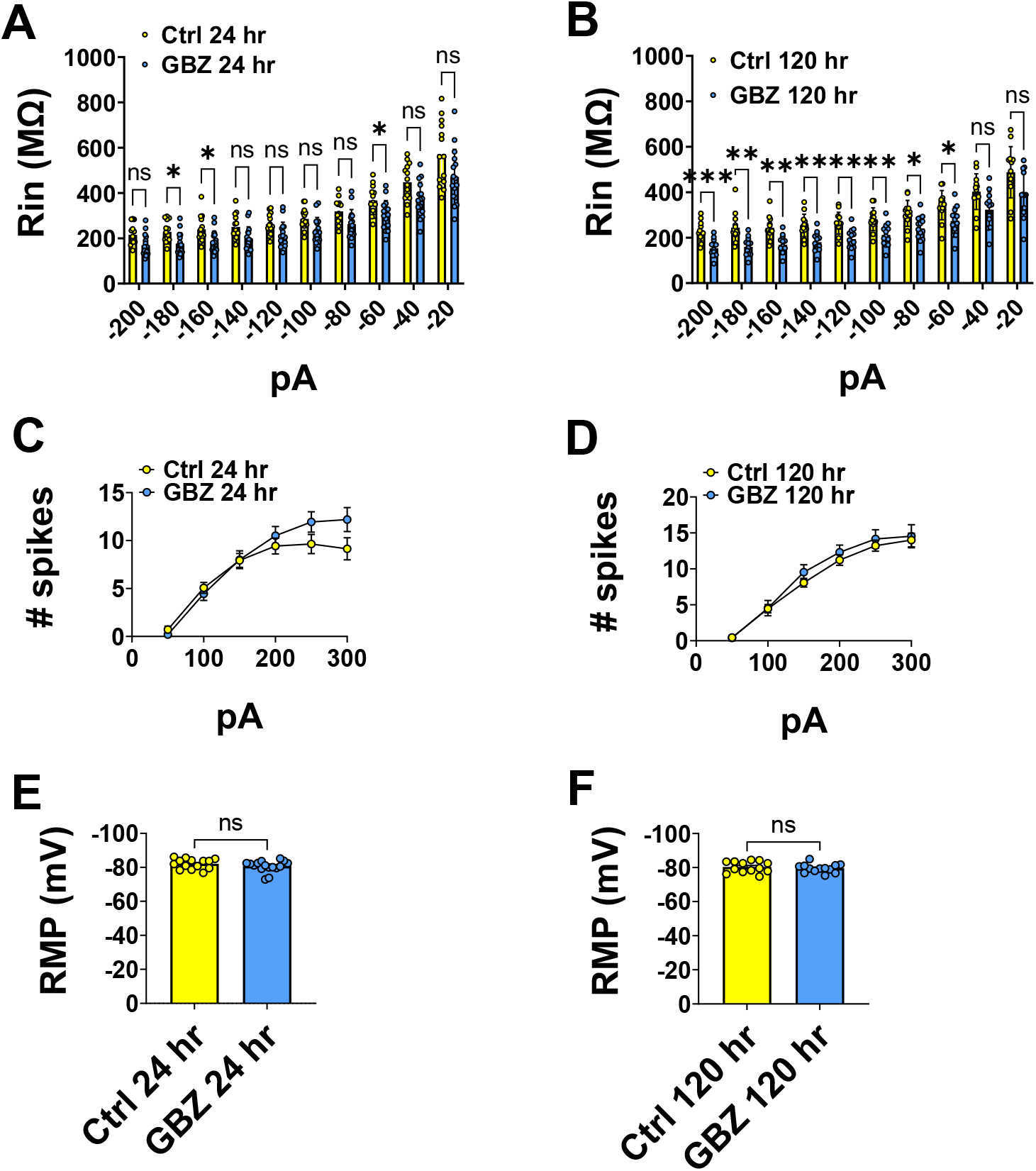
Effects of gabazine treatment on intrinsic measures. **A** and **B**: Modest reductions in R_in_ after 24 and 120 hrs of gabazine treatment (24 hr: Control vs. GBZ, main effect of group, *p <* 0.01; 120 hr: Control vs. GBZ, main effect of group, *p <* 0.001). **C** and **D**: no effect of gabazine treatment on evoked cell firing at 24 hrs or 120 hrs. **E** and **F**: no effect of gabazine on RMP at 24 hrs or 120 hrs. (n/group: 24 hrs: Control = 14, Gabazine = 16; 120 hrs: Control = 13, Gabazine = 13)

Gabazine has been used by others to induce downward homeostatic plasticity of firing rates and synaptic transmission (Goold & Nicoll, 2010; Vertkin et al., 2015). However, much of the inhibition onto GCs is tonic rather than phasic, and there is debate over gabazine’s efficacy in blocking this mode of inhibiting dentate GCs (compare Stell and Mody (2002) with Wlodarczyk et al. (2013)). Thus, we next tested the actions of a second GABA_A_ blocker, picrotoxin (PTX; 100 µM), which is regularly used to block phasic *and* tonic inhibition (Bai et al., 2001; Wlodarczyk et al., 2013), on mEPSCs in GCs. As with gabazine, we found no downscaling after 120 hr of PTX treatment and instead saw an increase in mEPSC frequency (**Fig. 3**). As a key control, we tested the effects of prolonged PTX treatment in the neighbouring CA3 region. Consistent with previous reports (Lee et al., 2013; Mao et al., 2018), we found robust homeostatic downregulation of mEPSC amplitude in CA3 pyramidal cells after 120 hr of PTX treatment, whereas frequency did not differ significantly between groups (**Fig. 4**).

**Figure 3:**
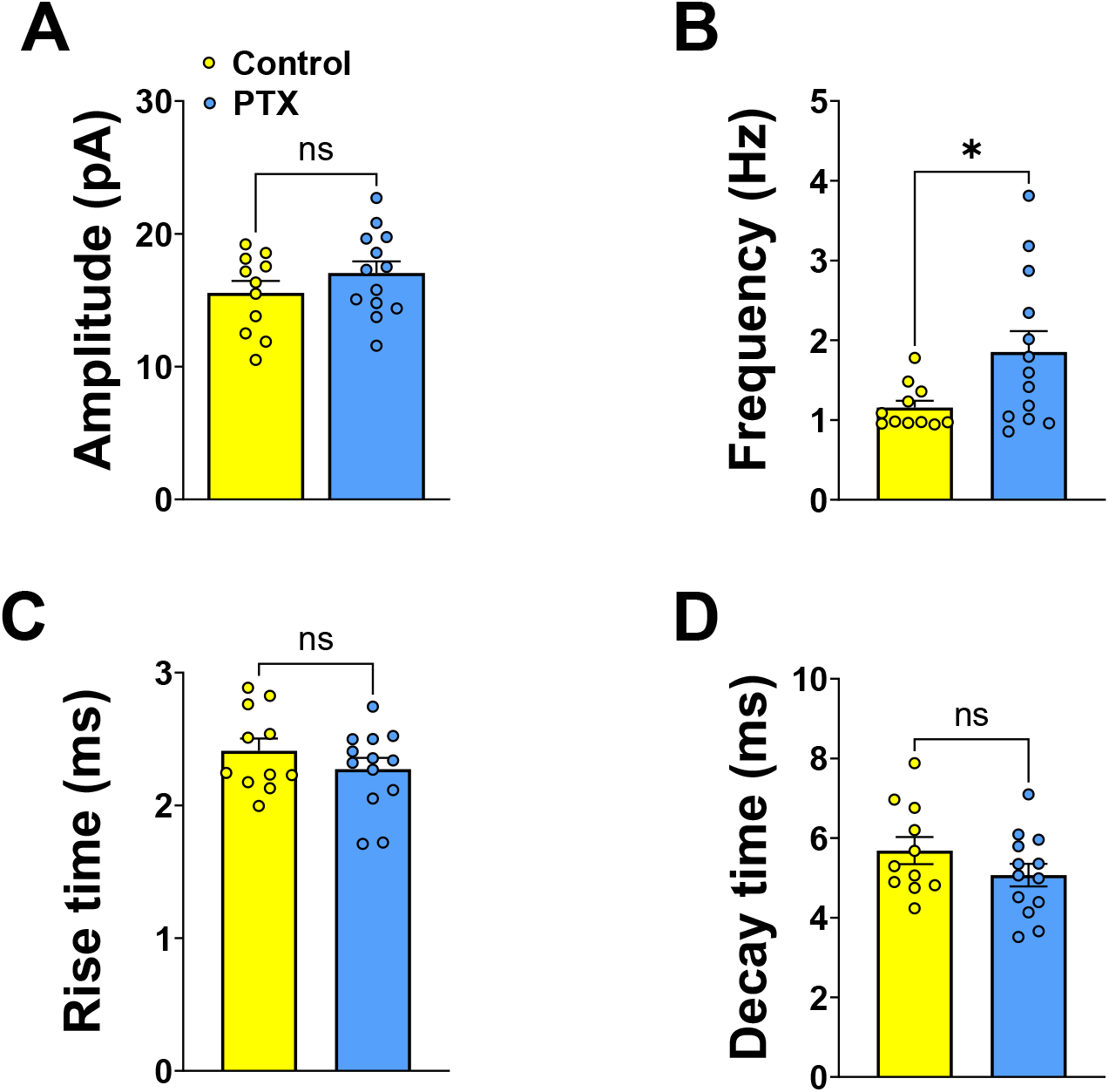
120 hr PTX treatment does not induce downscaling in dentate gyrus granule cells. **A:** mEPSC amplitude does not differ between Control a PTX treated cells at 120 hrs. **B:** mEPSC frequency is significantly increased after 120 hrs of PTX (Control: n = 11, 1.16 ± 0.1 Hz; PTX: n = 13, 1.85 ± 0.3 Hz, *p* = 0.029). **C** and **D:** no differences in mEPSC kinetics after 120 hrs of PTX.

**Figure 4:**
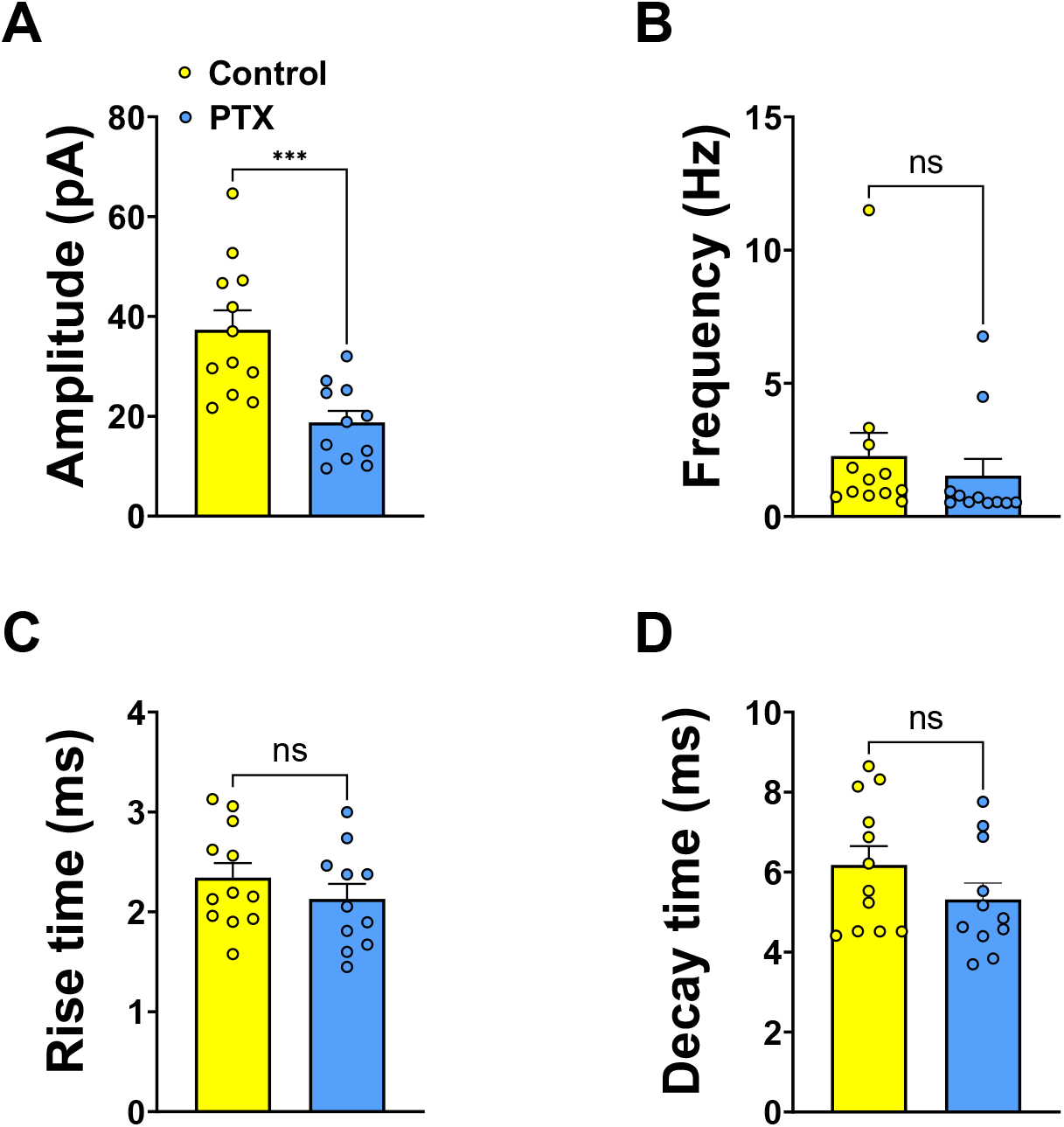
Downscaling of mEPSCs is evident in CA3. **A:** 120 hrs of PTX treatment elicits a significant reduction in mEPSC amplitude in CA3 (Control: n = 12, 37.3 ± 4 pA; PTX: n = 11, 18.8 ± 2 pA, *p* < 0.001). **B:** No effect of PTX on mEPSC frequency in CA3. **C** and **D:** No effect of PTX on mEPSC kinetics in CA3.

### Enhancing treatment duration and efficacy does not reveal synaptic downregulation

We next considered two possibilities that might account for an absence of downward homeostatic plasticity. Namely, the duration and efficacy of the treatment. We reasoned that the hyperpolarised resting V_m_ of granule cells (> -80 mV; Spruston & Johnston, 1992; Staley, Otis, & Mody, 1992), which is negative with respect to *E*_Cl-_ in GCs (Staley & Mody, 1992), may mean GABAergic antagonists in fact have a hyperpolarising effect (Chiang et al., 2012). Thus, GABAergic blockers alone may not greatly enhance firing in these cells (Chattipakorn & McMahon, 2003). To ensure augmented firing, we treated cultures with PTX and an additional 4 mM KCl. In acute application experiments on DIV 14 cultures, we verified this treatment elicited robust increases in cell firing (**Supplementary Fig. 1**). To address duration, we decided to prolong PTX & KCl treatment to a period beyond anything that might reasonably be considered necessary to produce a homeostatic response. To this end, slice cultures were maintained in control or PTX & KCl-supplemented media for 480 hr (20 days). Even after this extreme treatment, we found mEPSC amplitudes did not differ between groups. While we did not see the enhanced mEPSC frequency seen after gabazine or PTX alone, we also did not see a reduction of mEPSC frequency suggestive of homeostatic synaptic downregulation (**Fig. 5**).

**Figure 5:**
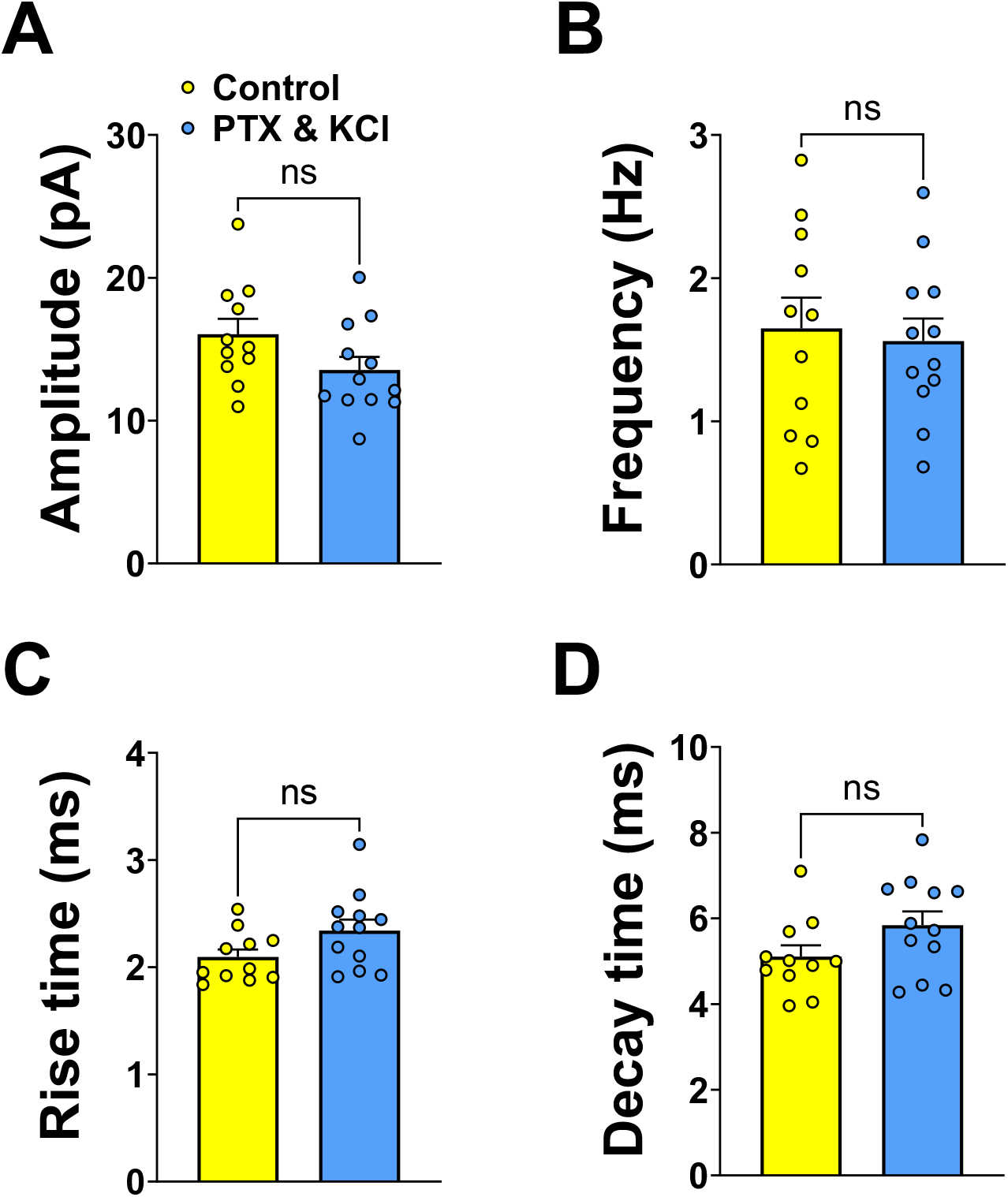
Downscaling is not evident after 20 days of PTX & KCl treatment. **A** and **B:** No effect of PTX & KCl on mEPSC amplitude or frequency, respectively, after 20 days. **C** and **D:** No effect of 20 day PTX & KCl treatment on mEPSC kinetics. (n/group: Control = 11, PTX & KCl = 12)

### Hebbian plasticity and synaptogenesis

Homeostatic synaptic plasticity is proposed to act as a negative feedback mechanism that counteracts activity changes induced by learning-related (i.e. Hebbian) forms of plasticity such as LTP and LTD (Abbott & Nelson, 2000; Turrigiano, 2017; Turrigiano & Nelson, 2000). However, we reasoned that ongoing Hebbian plasticity might be facilitated by GABA_A_ blockade (Wigstrom & Gustafsson, 1983, 1985), and that this may mask or counteract homeostatic changes. As these forms of plasticity are predominantly driven by NMDA receptor activation (Bliss & Collingridge, 2013), we incubated slices in the NMDA receptor blocker APV (50 µM) concurrently with PTX and KCl. Still, with NMDAR function blocked, we saw no effect of treatment on mEPSC amplitude and a robust potentiation of mEPSC frequency after 120 hr of treatment (**Fig. 6**).

**Figure 6:**
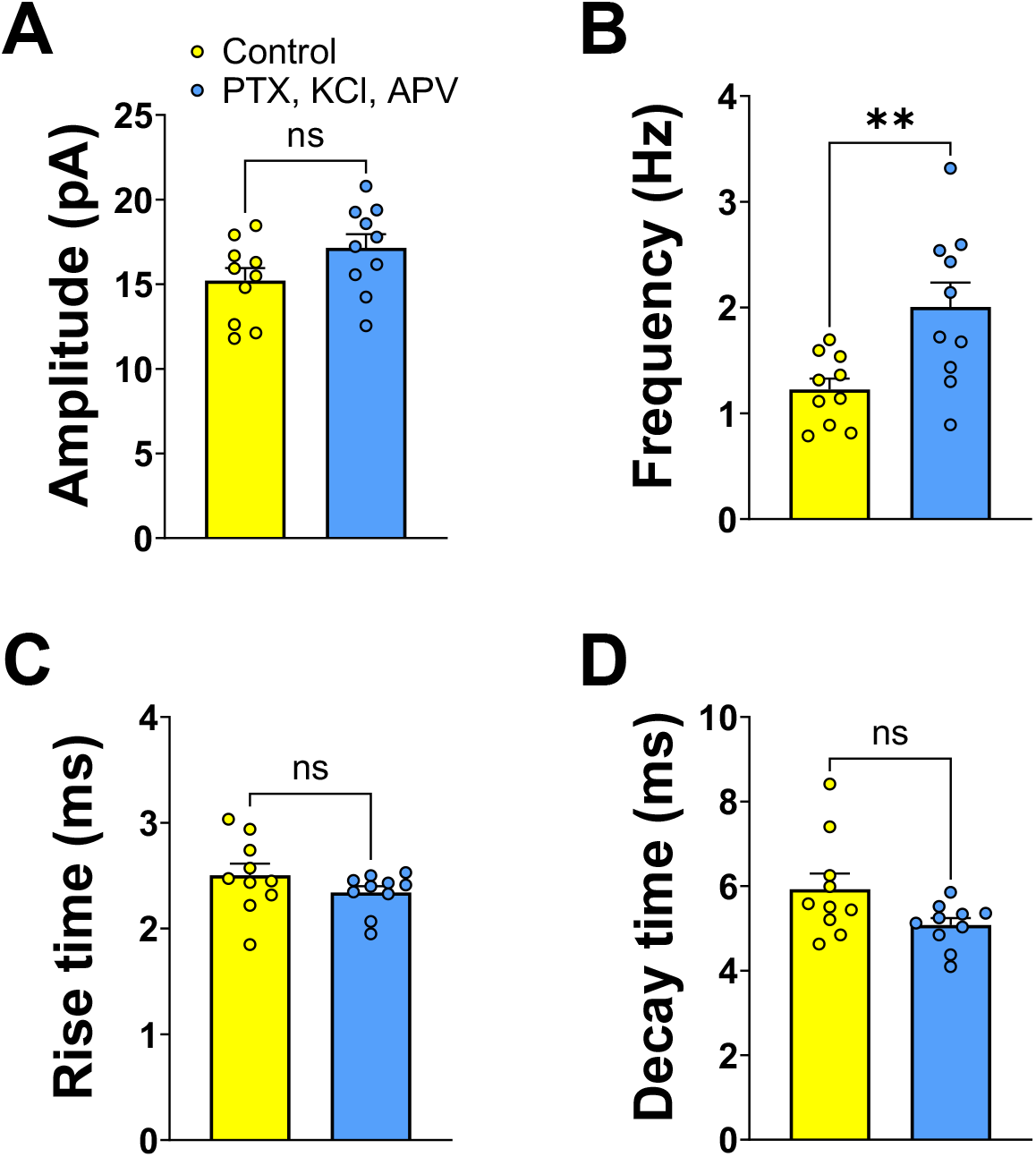
NMDAR blockade does not reveal synaptic downscaling. **A:** No difference in mEPSC amplitude between control cells and cells treated with PTX, KCl & APV for 120 hrs. **B:** mEPSC frequency is increased in cells treated with PTX, KCl & APV (Control: n = 10, 1.22 ± 0.1 Hz; PTX, KCL, APV: n = 10, 2 ± 0.2 Hz, *p* = 0.027). **C** and **D:** no effect of PTX, KCl & APV treatment on mEPSC kinetics.

Aside from Hebbian plasticity such as LTP and LTD, overall synaptic activity can be modulated through activity-dependent synaptogenesis. This is particularly relevant in the dentate gyrus, in which hyperactivated mossy fibres can sprout new collaterals that invade the molecular layer and form recurrent excitatory synapses onto neighbouring granule cells (Buckmaster, Zhang, & Yamawaki, 2002). We therefore reasoned that 5 days of PTX treatment might be triggering *de novo* synaptogenesis via mossy fibre sprouting, which would account for the elevated mEPSC frequency we observed and, potentially, mask downward homeostatic synaptic plasticity effects. Mossy fibre sprouting is triggered by L-type voltage gated calcium channels (Ikegaya, Nishiyama, & Matsuki, 2000), and *de novo* synaptogenesis is driven by astrocyte-derived thrombospondins (Eroglu et al., 2009). Thus, we repeated our PTX experiments but with concurrent application of gabapentin (100 μM), which blocks both L-type channels (Sutton, Martin, Pinnock, Lee, & Scott, 2002) and the thrombospondin receptor α2δ-1 (Field et al., 2006). Despite both signaling cascades being blocked, in these experiments we still saw greater mEPSC frequency after 120 hr of treatment (**Fig. 7**).

**Figure 7:**
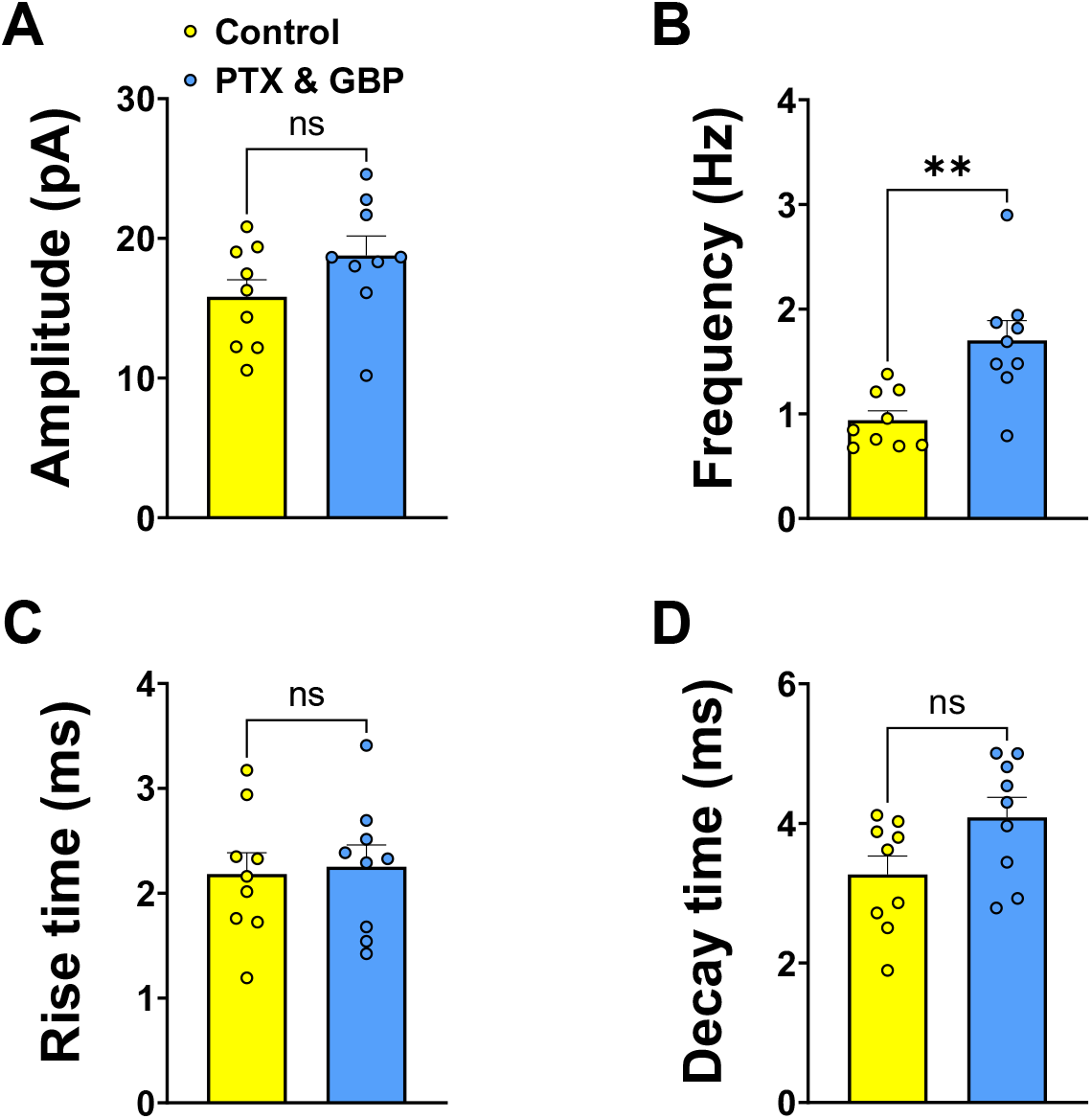
Thrombospondin blockade does not reveal synaptic downscaling. **A:** mEPSC amplitude does not differ between control cells and PTX & GBP treated cells after 120 hrs of treatment. **B:** mEPSC frequency is significantly greater in PTX & GBP treated cells (Control: n = 9, 0.94 ± 0.1 Hz; GBP: 1.7 ± 0.2 Hz, *p* = 0.004). **C** and **D:** No effect of treatment on mEPSC kinetics.

### Metaplasticity and synaptic homeostasis

Having failed to observe homeostatic synaptic downregulation in any of our experiments, despite the clear efficacy of our treatment in eliciting such plasticity in the neighbouring CA3 region, we were forced to conclude that it is highly unlikely that granule cells are capable of demonstrating overt homeostatic synaptic downregulation. We therefore pondered whether regulation at synapses onto granule cells might come through other means, following chronically elevated activity. In a previous study (Abraham, Mason-Parker, Bear, Webb, & Tate, 2001), it was reported that prolonged trains of backpropagating action potentials could metaplastically raise the threshold for LTP induction an PP-DG synapses. Metaplasticity is capable of maintaining synapse strength within a dynamic range by promoting or inhibiting Hebbian plasticity (Hulme et al., 2013). As the Bienenstock, Cooper and Munro model of metaplasticity posits an action potential driven shift in LTP/LTD thresholds (Bienenstock, Cooper, & Munro, 1982), we posited that chronically elevating activity might elicit similar changes to those seen during the acute (albeit strong) stimulation protocol employed by Abraham et al. To this end, we turned to an *in vivo* model in which mice were treated with PTX for 3 d prior to preparation of brain slices for LTP recordings. Animals were monitored for 1 hr post-injection, which is well beyond the reported half-life (15 min) of PTX when injected i.p. (Pressly et al., 2020). This dosage was subconvulsant, in accordance with previous reports (Ito, Lim, Nabeshima, & Ho, 1989; Nutt, Cowen, Batts, Grahame-Smith, & Green, 1982; Yokoro, Pesquero, Turchetti-Maia, Francischi, & Tatsuo, 2001). Using this protocol, which was by necessity milder than our *in vitro* treatments, we did not observe any changes in the average slope of field excitatory postsynaptic potentials (fEPSPs) generated at medial perforant path synapses with our test stimulation intensity, not did we see differences in the paired-pulse ratio to suggest any differences in transmitter release probability. However, we did see a significant reduction in the amount of LTP generated at these synapses (**Fig. 8**).

**Figure 8.**
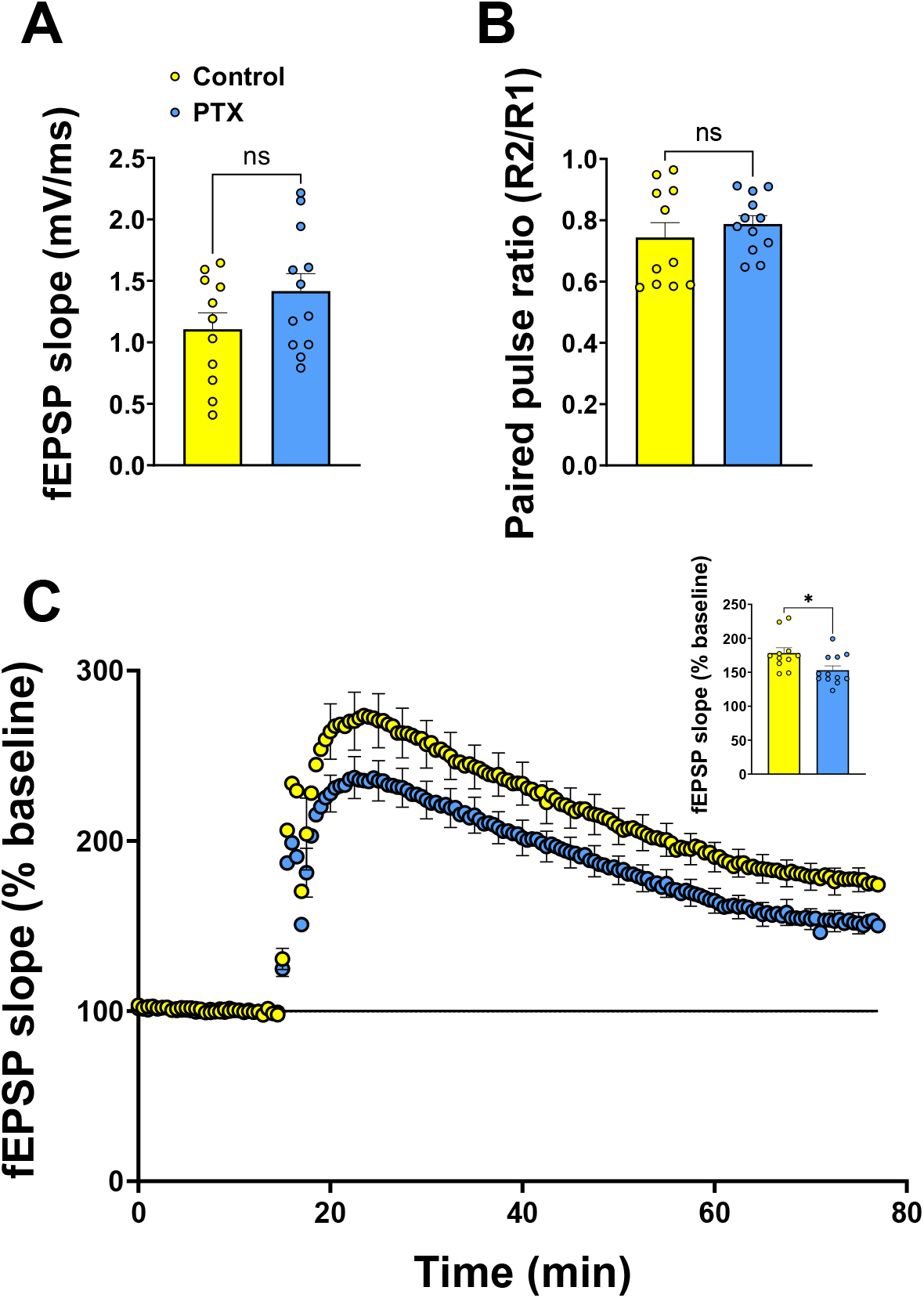
PTX treatment *in vivo* metaplastically inhibits LTP at medial perforant path synapses. **A:** No significant differences in basal fEPSP slope at test pulse intensity between slices from Control and PTX-treated animals. **B:** No significant differences in Paired Pulse Ratio between groups. **C:** Slices from PTX-treated animals exhibit a significant reduction in LTP (Control: n = 11, 178 ± 8%; PTX: n = 12, 153 ± 6%, *p* = 0.021). Inset: summary histogram of % change from baseline in final 10 min of recording.

## Discussion

We found no evidence of overt homeostatic downregulation of excitatory synaptic transmission onto granule cells in any of our attempts to elicit such plasticity. Instead, we saw only *enhanced* synaptic drive onto granule cells following chronic GABA_A_R blockade. To our knowledge, this is the first demonstration of a cell type that does not show downward homeostatic synaptic plasticity.

Why is downscaling absent at synapses onto granule cells? Certainly L-VGCCs, the purported triggers, are present on these cells (Hell et al., 1993). So too are the main molecular players identified elsewhere (Fernandes & Carvalho, 2016; Siddoway et al., 2014), such as the kinases CaMKIV (H. Wu et al., 2022) and polo-like kinase 2 (Kauselmann et al., 1999), the phosphatase PP1 (Sakagami, Ebina, & Kondo, 1994) and the immediate early gene Homer1a (Hoang, Böge, & Manahan-Vaughan, 2021). One intriguing exception here is the chromatin reader lethal 3 malignant brain tumor-like 1, which is expressed only weakly in granule cells but much more strongly in CA3 pyramidal neurons, where it is essential for downscaling (Mao et al., 2018).

Another notable distinction between synapses onto granule cells and those onto CA3 pyramidal cells is the expression of GABA_B_ receptors (GABA_B_Rs), which are a requirement for homeostatic downscaling (Vertkin et al., 2015). These receptors are either of the presynaptic GABA_B(1a)_ subtype or postsynaptic GABA_B(1b)_ subtype (Gassmann & Bettler, 2012). mRNA for either subtype is 30-50% lower in DG than in CA1 or CA3 (Bischoff et al., 1999), and shows highly laminar expression with GABA_B(1a)_ dominant at MPP synapses and GABA_B(1b)_ dominant at LPP synapses (Foster, Kitchen, Bettler, & Chen, 2013). Of note, both subtypes are present at MF-CA3 synapses (Kulik et al., 2003). It is therefore possible that downscaling is confined to synapses expressing both pre-*and* postsynaptic GABA_B_Rs.

Without downscaling of synapses or excitability, how do granule cells maintain their relative quiescence? Many mechanisms already work to maintain low firing rates in these cells, such as a highly hyperpolarised V_m_ (Staley et al., 1992) and substantial levels of tonic inhibition (Nusser & Mody, 2002). Moreover, although they may not trigger downscaling, L-VGCCs still fulfil homeostatic roles in granule cells by other means. Mendez and colleagues (2018) showed that spike trains in granule cells trigger a lasting reduction in spine formation, and the induction of LTP at MPP synapses triggers an L-VGCC-dependent heterosynaptic LTD at neighbouring LPP synapses (Christie & Abraham, 1994). Considering these phenomena alongside the metaplastic mechanisms seen here and elsewhere (Abraham et al., 2001), granule cells appear sufficiently well-armed to maintain the quiescence required for their purported physiological roles. Indeed, it has previously been noted that granule cells are, in the main, biased towards exhibiting synaptic depression (Lisman, 2011), which may be why this largely silent pool of cells requires ongoing replenishment through neurogenesis. The absence of homeostatic downscaling would therefore only become problematic following pathologically elevated activity such as seizure. In this sense, homeostatic downscaling may be better couched as a response to pathologically high activity, rather than responses to learning-related changes as has been suggested (Abbott & Nelson, 2000; Keck et al., 2017; but see: C. H. Wu et al., 2021).

What consequence does the absence of downscaling have for granule cells? The dentate gyrus is profoundly affected in temporal lobe epilepsy, making it tempting to speculate that the lack of downscaling contributes to the susceptibility of this region. While experimental status epilepticus in rats has been shown to increase miniature inhibitory postsynaptic currents in granule cells (Leroy, Poisbeau, Keller, & Nehlig, 2004), inhibitory interneurons are frequently compromised or lost in TLE models (Sloviter, 1991; Sun, Mtchedlishvili, Bertram, Erisir, & Kapur, 2007). Thus, after prolonged status epilepticus there may be no remaining mode of homeostatic plasticity that can sufficiently regulate local DG circuits. On the note of TLE, we should contrast the upregulated synaptic activity in our experiments with the phenomenon of mossy fiber sprouting (T. Sutula, Cascino, Cavazos, Parada, & Ramirez, 1989; T Sutula, He, Cavazos, & Scott, 1988), in which GC axons reorganise post-status epilepticus to extend into their own dendritic fields and create recurrent excitatory loops. While it would be tempting to suggest we are seeing the same effect, we note that the two reported triggers of sprouting, NMDARs and L-VGCCs (Ikegaya et al., 2000; T. Sutula, Koch, Golarai, Watanabe, & McNamara, 1996) do not contribute to the increased mEPSC frequency we saw. Thus, it appears that a second, mechanistically distinct mode of increased excitatory drive onto granule cells is seen following prolonged hyperactivity. We conclude that granule cells of the DG are therefore well-placed to remain quiescent under normal physiological conditions but are poorly equipped to adapt to chronic elevations in activity.

An outstanding matter is the mechanism by which synaptic transmission is upregulated in granule cells after chronic depolarisation. Having already ruled out classic Hebbian LTP, mossy fibre sprouting and *de novo* synaptogenesis via thrombospondins, there are few options remaining. Naturally, other triggers of synaptogenesis such as BDNF could be involved (Cunha, Brambilla, & Thomas, 2010). Another possibility is increased release probability. Although this was not evident from paired-pulse testing in our acute slices, these experiments only tested one input pathway to the DG (and are in any case different from our organotypic slice experiments). For example, our treatments may cause sprouting or increased glutamate release from mossy cells that typically innervate the inner molecular layer (Scharfman, 2016). However, this may be unlikely as innervation of granule cells by mossy cells is in fact reduced after status epilepticus (Butler, Westbrook, & Schnell, 2022). Thus, the mechanism(s) underlying enhanced transmission onto granule cells remain enigmatic and a focus for future investigations.

## Acknowledgements

We would like to thank the lab of Thomas Deller for assistance with optimising the preparation of organotypic slice cultures. This work was funded by a Brain Research New Zealand Fellowship, a University of Otago Research Grant, and a Maurice and Phyllis Paykel Trust Project Grant to ODJ.

## Author contributions

ODJ conducted the bulk of experimental work and data analysis. KB contributed to patch clamp recordings and data analysis. ODJ, RME and WCA designed experiments and wrote the manuscript.

---

**Supplementary Figure 1:**
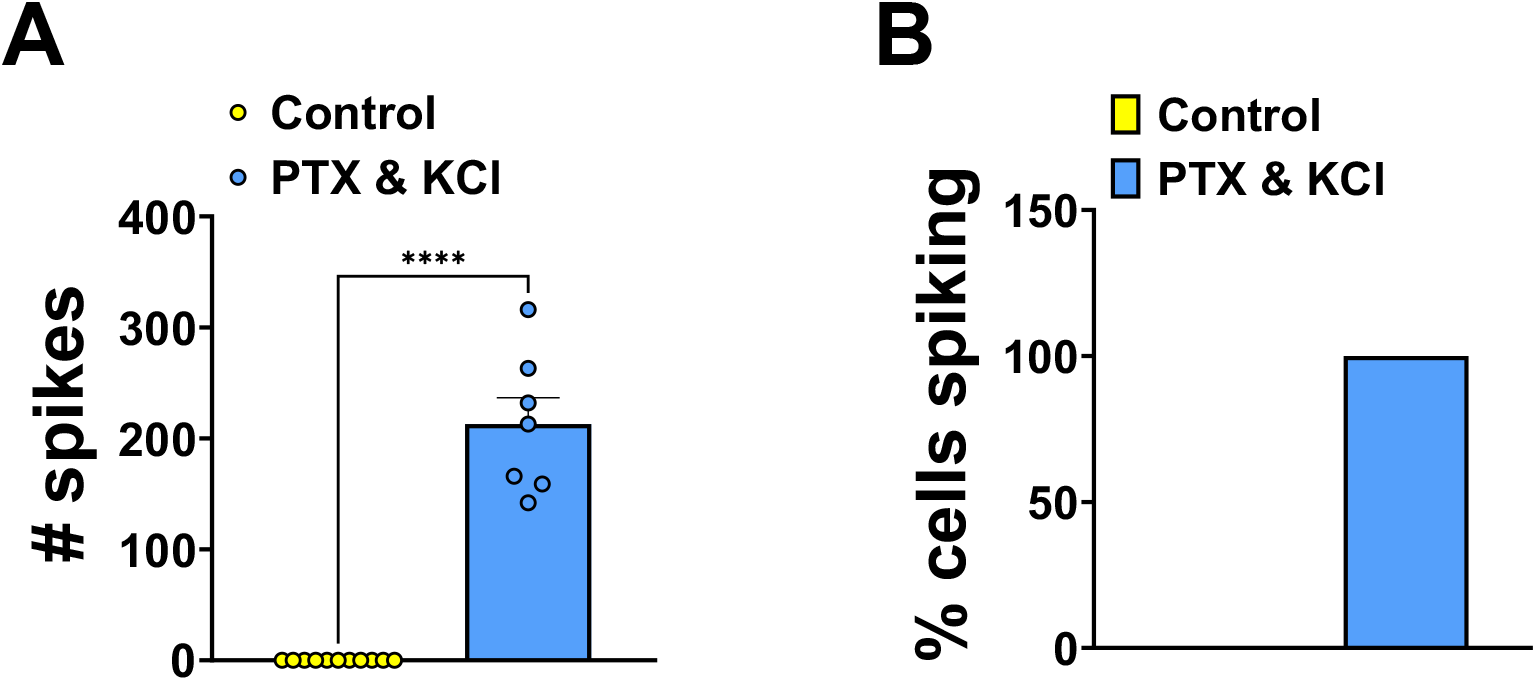
Validation of spiking in presence of PTX & KCl. **A:** Combined PTX & KCl treatment led to a significant increase in cell firing during a 10 min application period (Control: n = 10, 0 ± 0 spikes; PTX & KCl: n = 6, 213 ± 15 spikes, *p*, 0.0001). **B:** Cell firing was not seen in any control cells but was seen in 100% of cells exposed to PTX & KCl.

